# DsbA is a redox-switchable mechanical chaperone

**DOI:** 10.1101/310169

**Authors:** Edward C. Eckels, Deep Chaudhuri, Soham Chakraborty, Daniel J. Echelman, Shubhasis Haldar

## Abstract

DsbA is a ubiquitous bacterial oxidoreductase that associates with substrates during and after translocation, yet its involvement in protein folding and translocation remains an open question. Here we demonstrate a redox-controlled chaperone activity of DsbA, on both cysteine-containing and cysteine-free substrate, using a magnetic tweezers-based single molecule force spectroscopy that enables independent measurements of oxidoreductase activity and chaperone behavior. Interestingly we found, this chaperone activity is tuned by the oxidation state of DsbA; oxidized DsbA is a strong promoter of folding, but the effect is weakened by reduction of the catalytic CXXC motif. We further localize the chaperone binding site of DsbA using a seven-residue peptide which effectively blocks the chaperone activity. We calculated that DsbA assisted folding of proteins in the periplasm generates enough mechanical work to decrease the ATP consumption needed for periplasmic translocation by up to 33%. In turn, pharmacologic inhibition of this chaperone activity may open up a new class of anti-virulence agents.

## Introduction

Peptide translocation is of utmost importance in bacteria, where a large number of proteins, including virulence factors, must pass through the translocon pore in the unfolded state and properly fold into the native configuration in the periplasm(*1*–*5*). A large number of proteins secreted into the periplasmic space contain disulfide bonds which are introduced by oxidoreductase enzymes of the Dsb family that share a conserved thioredoxin-type fold(*6*, *7*). The prototypical family member, DsbA, is necessary for the maturation of a range of virulence factors including flagellar motors and pilus adhesins, Type III secretion systems, and heat-labile and heat-stable enterotoxins and has emerged as a target for novel antibiotic development(*8*–*11*). Recent studies shows DsbA assist efficient secretion of cysteine-null proteins, indicating its ability to engage unfolded substrates during translocation (*14*).

The well-studied eukaryotic chaperones BiP and Hsp70 are known to facilitate translocation of peptides into the endoplasmic reticulum and mitochondria, respectively(*15*–*17*). It is believed that chaperone binding to an unfolded substrate protein on one side of the membrane can induce biased transport of a polypeptide through a Brownian ratchet mechanism(*18*–*21*). This relies not on the energy of hydrolysis of ATP but on the differential concentrations of the chaperones on either side of the membrane. Simple binding of chaperones on one side of the membrane prevents backsliding of the protein into the pore, providing directional transport of the polypeptide. Once the polypeptide is properly folded on the other side of the membrane, it is too large to pass back through the pore. Although DsbA is considered to be a ‘weak’ chaperone, the role of oxidation on its chaperone activity or periplasmic translocation is not known (12, 13). If DsbA can indeed redox-dependent chaperone under the physiological constrain, faced by the protein in translocon pore, it may play a much broader role in periplasmic secretion than previously recognized.

Peptides emerging from nanoscale tunnels such as the translocon pore are known to experience an effective stretching force due to molecular confinement(*22*–*24*). As such, these peptides emerge from the pore in a linear, extended state and can either collapse on the mouth of the pore or engage with soluble chaperones in the periplasm. Here we apply single molecule magnetic tweezers assay to mimic the entropic stretching forces of the translocon pore. Using this assay, soluble DsbA is shown to cleave and reform disulfide bonds in a model immunoglobulin protein domain, validating the chaperone activity of the oxidoreductase enzyme. We further investigate the redox dependent chaperone activity of DsbA with cysteine free substrates using the globular B1 domain of protein L, which undergoes an equilibrium between the folded and unfolded states under small pulling forces (4-9 pN). We find that the presence of soluble DsbA in the experimental buffer greatly increases the residence time of protein L in the folded state and allows the protein L domain to refold at higher forces. Using the residence time of protein L in the folded state as our metric, we demonstrate the ability to turn DsbA chaperone activity “on” and “off” by changing the redox state of its active site disulfide bond or by targeting it’s putative binding site with a small inhibitory peptide. These results suggest that DsbA acts as a mechanical foldase that can engage and fold peptides under an effective stretching force, such as those emerging from the translocon pore. Because DsbA drives folding of its substrates at higher forces, it increases the amount of mechanical work performed by the folding substrate, which in turn decreases the energy needed to liberate it from the pore. We calculate that the work done by chaperone assisted protein folding on the periplasmic side of the translocon pore could be an energy source that greatly decreases the ATP consumption of the various translocation motors.

## Results

### Monitoring single molecule oxidative folding by DsbA

We recently developed a single molecule assay to study chaperone activity using magnetic tweezers-based force spectroscopy that overcomes limitations of bulk studies of chaperones(*25–28*). By their nature, bulk studies examine chaperone-client interactions in an environment far from *in vivo*, and denaturants such as guanidine, low pH, or high temperature used in these experiments necessarily perturb the structure of the chaperone as well as the substrate protein. Single molecule magnetic tweezers have the unique advantage of using mechanical force to denature only the substrate protein without disturbing chaperone molecules in the surrounding solution. Mechanical unfolding of individual proteins is achieved through attachment of the substrate protein to microscopic probes; in our magnetic tweezers-based force spectroscopy, proteins are tethered at their N-terminus to a glass surface via HaloTag-Halo ligand covalent chemistry(*27*, *29*), and at their C-terminus to a paramagnetic bead via biotin-streptavidin(*30*) (Fig 1A, inset). Positioning of a permanent magnet at fixed distances from the paramagnetic bead applies a passive force clamp that can explore a broad range of forces up to ~100 pN. Furthermore, the long-term stability of magnetic tweezers enables measurement of slow rates, with successive measurements from a single protein achieving a record of two weeks(*27*).

**Figure 1:**
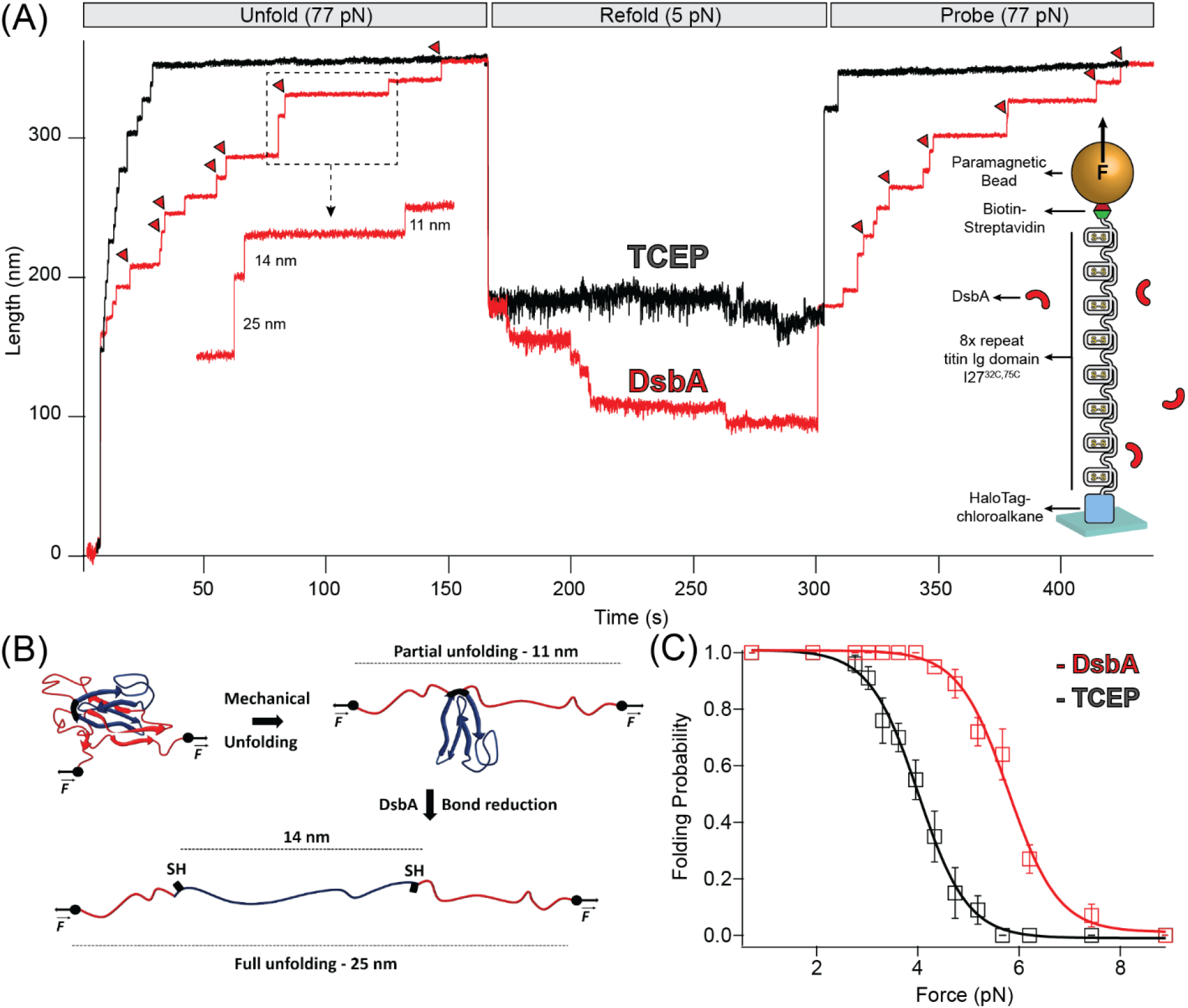
Chaperone activity of DsbA monitored in disulfide bonded protein: (A) Reduction of I27 by TCEP and DsbA: In presence of reducing condition maintained by 10 mM TCEP (black trace), the polyprotein is unfolded at denature pulse of 77 pN, causing a mixture of six 11 (due to mechanical unfolding) and 14 nm (due to disulfide reduction) along with two already reduced 25 nm steps. After unfolding, the force is quenched to 5.2 pN for observation of refolding events. A final probe pulse at 77 pN reveals only a single 25 nm step, representing the refolding of a single I27 domain in its reduced state. With DsbA (red trace), unfolding at denature pulse at 77 pN results in similar seven 11 and 14 nm steps (red triangles indicating enzymatic disulfide reduction) with a single 25 nm step, but after quenching the force to 5.2 pN, seven downward folding steps occurred, in contrast to only one folding step in presence of TCEP. A final probe pulse shows six mixed steps of 11 and 14 nm and one 25 nm steps, indicating re-oxidation of six Ig domains. **(B) Schematics of complete unfolding of a titin Ig domain under force:** Application of mechanical force on I27^C32-C75^ first unfolds the protein up to the internal disulfide bond, leading to an extension of 11 nm (of the un-sequestered peptide) that is followed by an additional extension of 14 nm due to reduction of the disulfide bond. **(C) Folding probability increased in presence of DsbA:** Folding probability as a function of force is plotted in presence of TCEP (black) and DsbA (red). The folding probability is shifted to a higher force in the presence of DsbA, with the force required for 50% refolding probability increasing from 4.1 to 5.8 pN. More than eight individual trajectories are measured and averaged for each force. More than five individual trajectories are measured and averaged for each force. Error bars are standard errors of mean (s.e.m.)

We used the magnetic tweezers assay to first measure the oxidoreductase activity of DsbA on a model titin immunoglobulin domain, I27^C32-C75^, containing a single disulfide bond(*31*, *32*). The recombinant protein construct was engineered with eight tandem repeats of the I27^C32-C75^ to provide a molecular fingerprint upon unfolding. Unfolding of an oxidized I27^C32-C75^ domain containing a formed disulfide bond results in an 11 nm extension(*25*, *31*) of the polypeptide chain (Fig 1A,B). Introduction of a reducing agent to the solution results in cleavage of the I27^C32-C75^ disulfide bond and release of 14nm of cryptic length from the polypeptide sequestered behind the disulfide bond (Fig 1A,B). In the assay, the eight tandem repeats are unfolded during the “denature” pulse at a pulling force of 77 pN in the presence of reduced 50 μM DsbA (Fig 1A, red trace) or 10 mM tris carboxyethyl phosphine (TCEP) (Fig 1A, black trace). The protein is left at this high force for sufficient time to allow for unfolding (11 nm steps) and reduction (14 nm steps) of all eight I27^C32-C75^ domains. The force is then “quenched” to 5.2 pN to allow for refolding for ~150 seconds, followed by a subsequent “probe” pulse back to 77pN. The probe pulse allows for the counting of the number of re-oxidized I27^C32-C75^ domains, which is measured as a function of the quench force. In the two recording shown in Fig 1B, seven I27^C32-C75^ domains refolded during the quench pulse in the presence of DsbA, whereas only one domain refolded in presence of reducing agent TCEP. Of the seven domains that refolded in the presence of DsbA, six contained reformed disulfide bonds (11 nm steps), while one refolded with its cysteines still reduced (25 nm step)

This assay was performed for many cycles, and Fig 1C summarizes the folding probability of I27^C32-C75^ in the presence and absence of DsbA. The folding probability, *P_f_*, describes the likelihood of a single domain to be folded at a particular force (see Supp. Fig 1 and Methods). The half-point force for I27^C32-C75^ (denoted as folding probability = 0.5) shows a prominent rightward shift from 4.1 to 5.8 pN in presence of DsbA. A similar shift in the folding probability of I27^C32-C75^ was also noted for protein disulfide isomerase (PDI), a eukaryotic oxidoreductase from the thioredoxin family(*25*). The folding step size noted during the quench pulse indicates that folding occurs before introduction of the disulfide bond(*33*, *34*), suggesting that there is acceleration of the folding process due to the chaperone is likely independent from conformational restriction by a newly introduced disulfide bond.

### DsbA chaperones a cysteine-free substrate in a redox dependent manner

The B1 antibody light chain binding domain of protein L (hereafter referred to as “protein L”) from *Finegoldia magna* is a model substrate well described in bulk biophysical and single-molecule force spectroscopy techniques(*35*–*38*). Protein L is 62 residues in length with a simple α/β fold (Fig 2B) that, importantly, lacks any metal cofactors, or cysteine or proline residues(*39*). In all studies herein, we employ an 8-repeat tandem modular protein L, flanked with N-terminal HaloTag and C-terminal biotin for tethering in the same geometry as used for I27^C32-C75^. With application of a high mechanical force (45 pN in Supp. Fig 1A, Pulse I), the eight protein L repeats unfold as eight discrete stepwise extensions of 15 nm, providing the distinct single-molecule fingerprint of the polyprotein (Fig 2A, inset). Upon a decrease in force, the total length of the fully unfolded polypeptide collapses due to polymer entropy, followed by discrete stepwise contractions from individual protein L domain refolding (9.0 nm; Supp Fig 1A, Pulse II). After several seconds, an equilibrium is reached between the stepwise extensions of unfolding and the stepwise contractions of refolding, with an identical length for each transition (Supp Fig 1A, Pulse II). The equilibrium behavior is reported by the folding probability *P_f_* as described previously(*26*) (Supp Fig 1A, also Methods).

**Figure 2.**
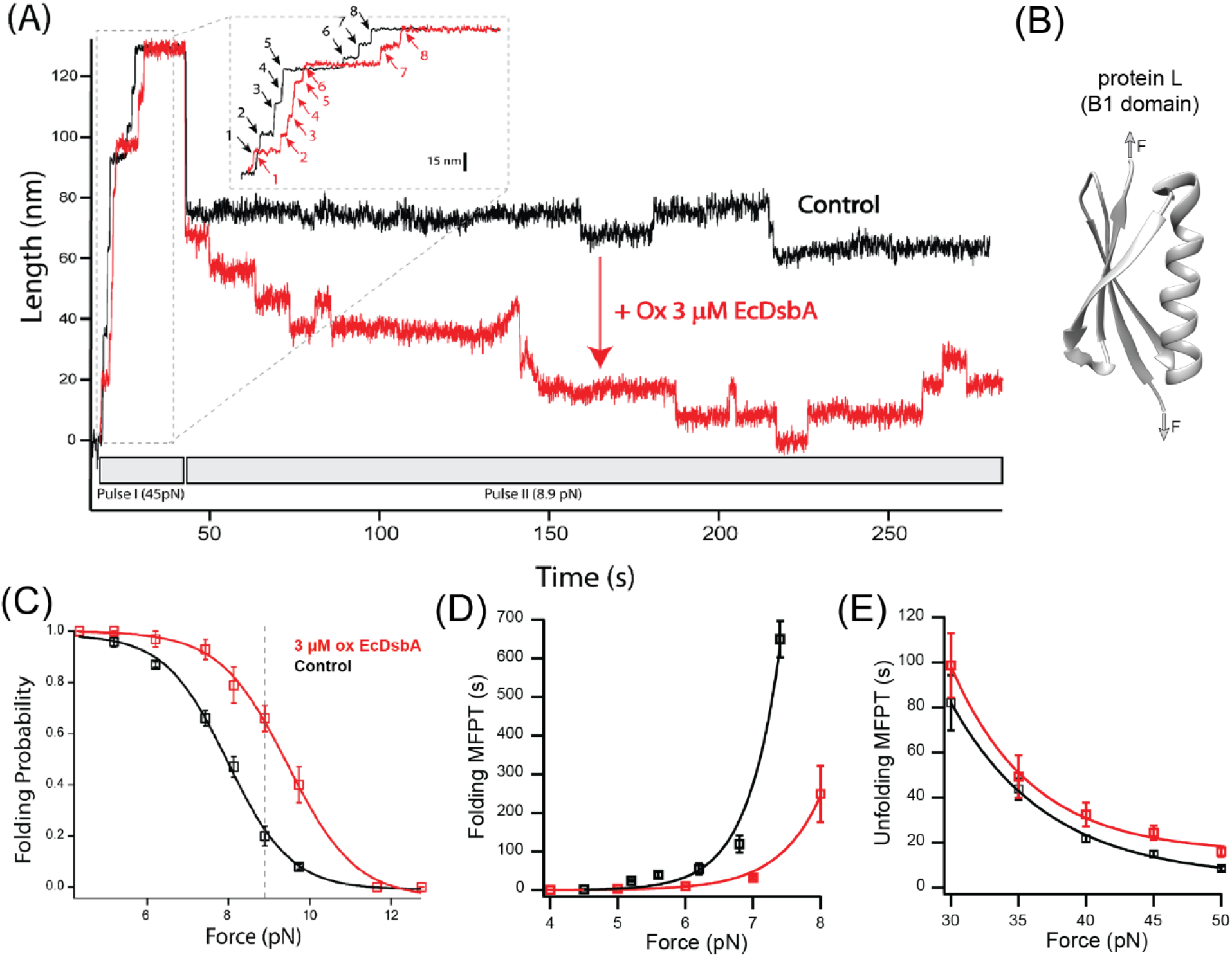
DsbA chaperones the protein L domain (A) Magnetic tweezers force-clamp trajectory of an protein L octamer in the presence (red) and absence (black) of 3 μM oxidized DsbA at 8.9 pN. In Pulse I, the protein is unfolded at 45 pN, where the 8 protein L unfolding events are observed as steps of ~15 nm (see inset for horizontally magnified trace). In Pulse II, force is quenched to 8.9 pN, where an initial relaxation refolding is observed as downward steps of ~9.5 nm, followed by equilibrium unfolding/refolding steps. **(B) B1 domain of protein L** showing a simple α/β fold without any cysteine or disulfide moieties. **(C) Folding probability as a function of force in the presence (red) and absence (black) of DsbA**. The rightward shift of the curve in the presence of DsbA indicates mechanical chaperone activity of DsbA that enhances folding of protein L in the force range from 4-11 pN. The maximum difference in the folding probabilities, marked by the gray line, occurs at 8.9 pN, the force demonstrated in panel (A). Error bars represent the standard errors of the mean of ≥ 5 independent molecules. **(D) Variation in MFPT of refolding:** The MFPT of refolding is plotted as a function of force in absence (black) and presence (red) of DsbA. The MFPT data were fit with a single exponential equation *MFPT(F)* = *Ae^-F/φ^* More than eight individual trajectories are measured and averaged for each force. Error bars are standard errors of mean **(E) Variation in MFPT of unfolding:** DsbA negligibly delays the unfolding kinetics by marginally increasing the time of complete unfolding of all eight domains. The difference in MFPT at the lowest unfolding forces (30 and 35 pN) is not statistically significant. More than eight individual trajectories and more than sixty unfolding events has been calculated for each force. Error bars are standard errors of mean (s.e.m.)

For protein L, the equilibrium position between folding and unfolding is sharply force-dependent (Fig 2C), with the folding probability shifting from >99% at 4 pN to <1% at 12 pN. In the absence of DsbA, the 50% folding probability is found at 8.0 pN. Upon addition of 3 μM of oxidized DsbA to the magnetic tweezers experiment, a shift in the folding probability is observed (Fig 2A). At an equilibrium force of 8.9 pN, the protein L polyprotein transitions between its 5th, 6th, and 7th folded state in the presence of oxidized DsbA (Fig 2A, red trace), whereas in the absence of oxidized DsbA the polyprotein transitions between the fully unfolded and the 1st folded state (Fig 2A, black trace). At the extremes of 4 pN and 12 pN, folding is unaffected, with either 100% of the substrate folded (at 4 pN) or unfolded (at 12 pN), independent of the presence of the enzyme. However, at intermediate forces, the effect of DsbA becomes more pronounced (Fig 2C). The largest shift is observed at 8.9 pN (dotted line, Fig 2C), where the protein L substrate has a 0.66 ± 0.05 probability of being folded when in the presence of oxidized DsbA, versus a 0.20 ± 0.04 probability of being folded in its absence.

### Effect of DsbA on mean first passage time (MFPT) of unfolding and refolding of protein L

We systematically explored the effect of DsbA by comparing the kinetics of protein L in both high and low force regime, with and without 3μM oxidized DsbA. The kinetic analysis allows one to determine if the shift in folding probability by DsbA is caused by a decrease of the energy of the folding transition state, an increase in the energy of the unfolding transition state, or both. The kinetics of unfolding and refolding can be characterized by mean first passage time (MFPT), where the first passage time (FPT) is denoted as the minimum time taken to complete unfolding and refolding of all eight domains. Mean first passage time (MFPT) is determined by averaging FPT over several trajectories(*26*). Figure 2D and 2E illustrate the comparison in the MFPT of refolding and unfolding in absence (black) and presence (red) of DsbA. In the presence of DsbA, the MFPT of refolding decreases at a particular force magnitude and thus, shows an accelerated folding kinetics by DsbA (Fig 2D). In contrast to refolding, DsbA marginally delays the unfolding kinetics (Fig 2E) at forces from 30-50 pN. Although the difference is small, it suggests that DsbA may be able to associate with the collapsed protein L substrate as well as the denatured form. Fit parameters for the single exponentials can be found in the supplementary information.

### Oxidation dependent activity of DsbA

Within the Gram-negative periplasm, oxidized DsbA represents the active fraction that is capable of forming mixed disulfides between enzyme and substrate peptide. After transferring the disulfide bond into its substrate, DsbA is released with both cysteines of its active site *CXXC* motif reduced. We therefore asked whether the chaperone-like activities of DsbA might depend on its redox state, which is expected if the chaperone activity originates from the hydrophobic groove encapsulating the catalytic site. We reduced freshly purified DsbA with an overnight incubation in 100 μM TCEP (which was then removed with a 10 kDa cutoff concentrating column) and repeated the magnetic tweezers-based mechanical foldase assay. Whereas 3 μM of oxidized DsbA shifts the probability of substrate folding to 0.66 at 8.9 pN, 3 μM of reduced DsbA (Supp Fig 2) induces no such shift in substrate folding (Fig 3A). Experiments with reduced DsbA tracked the folding probability of the protein L substrate alone over the range of 5 – 11 pN (Fig 3A). A comparable foldase effect with reduced DsbA requires 50 μM, a ~17-fold excess over oxidized DsbA (Fig 3B, green curve and Supp Fig 3). Importantly, these experiments with reduced DsbA were performed with 100 μM of reducing agent TCEP in solution to prevent spontaneous re-oxidation of the catalytic disulfide bond. These data demonstrate that the catalytic cysteines are likely part of the binding surface responsible for the chaperone activity of DsbA and that alteration of the redox state of the cysteines modulates substrate affinity, possibly by changing the charges or the hydrophobicity of the binding surface, or by altering the overall conformation of the enzyme(*40*, *41*).

**Figure 3:**
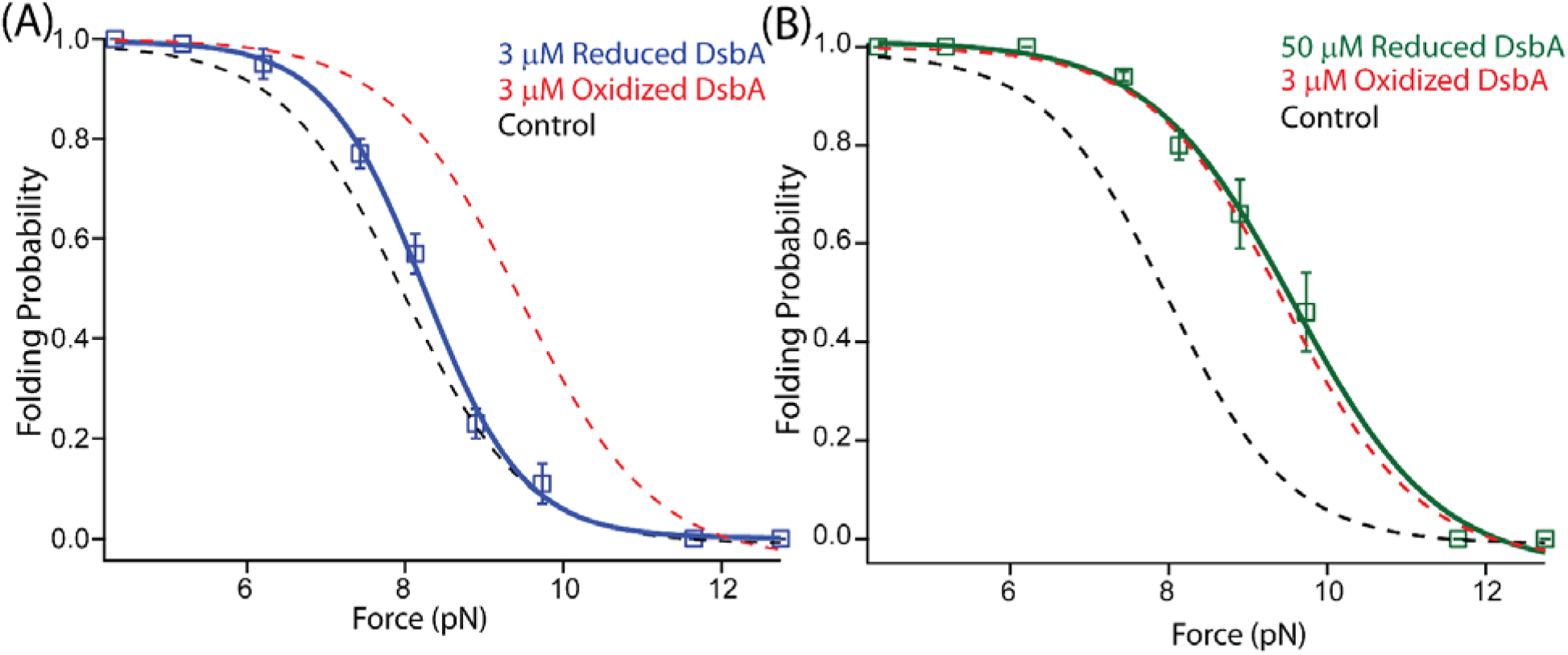
A redox switch controls the chaperone activity of DsbA: **(A)** The folding probability of 3 μM reduced DsbA (blue squares) has no significant effect on the folding probability of protein L, coinciding with the control experiment (black dotted line). More than five individual trajectories are measured and averaged for each force. Error bars are standard errors of mean **(B)** Increasing the concentration of reduced DsbA to 50 μM restores the mechanical chaperone activity of DsbA. The folding probability of protein L as a function of force in the presence of 50 μM reduced DsbA (green squares) coincides with the folding probability curve of the 3 μM oxidized DsbA (red dotted line). The fittings in the absence (black dotted line) and in the presence of 3 μM oxidized DsbA (red dotted line) is plotted as control. More than seven individual trajectories are measured and averaged for each force. Error bars are standard errors of mean.

### A peptide antagonist blocks the chaperone activities of DsbA

DsbA has emerged as an attractive therapeutic target due to its vital role in the maturation of bacterial virulence factors, including toxins, adhesins, and secretion machinery(*10*, *42*–*44*). For this reason, several peptide inhibitors of DsbA have been developed, based off the oxidoreductase interacting partner DsbB, which binds at and oxidizes the catalytic *CXXC* via interaction by a short loop with the sequence PFATCDF(*8*). A single point mutation of the N-terminal phenylalanine to tryptophan (and substitution of the N-terminal residue to serine to improve solubility) produced the peptide PWATCDS with high affinity for the hydrophobic groove of DsbA from *Proteus mirabilis* in a co-crystal structure(*9*) (Fig 4C). Binding of PWATCDS effectively blocks the oxidoreductase activity of DsbA. We therefore asked whether this peptide binding at the hydrophobic groove neighboring the *CXXC* motif would also inhibit the mechanical foldase activity of DsbA.

**Figure 4:**
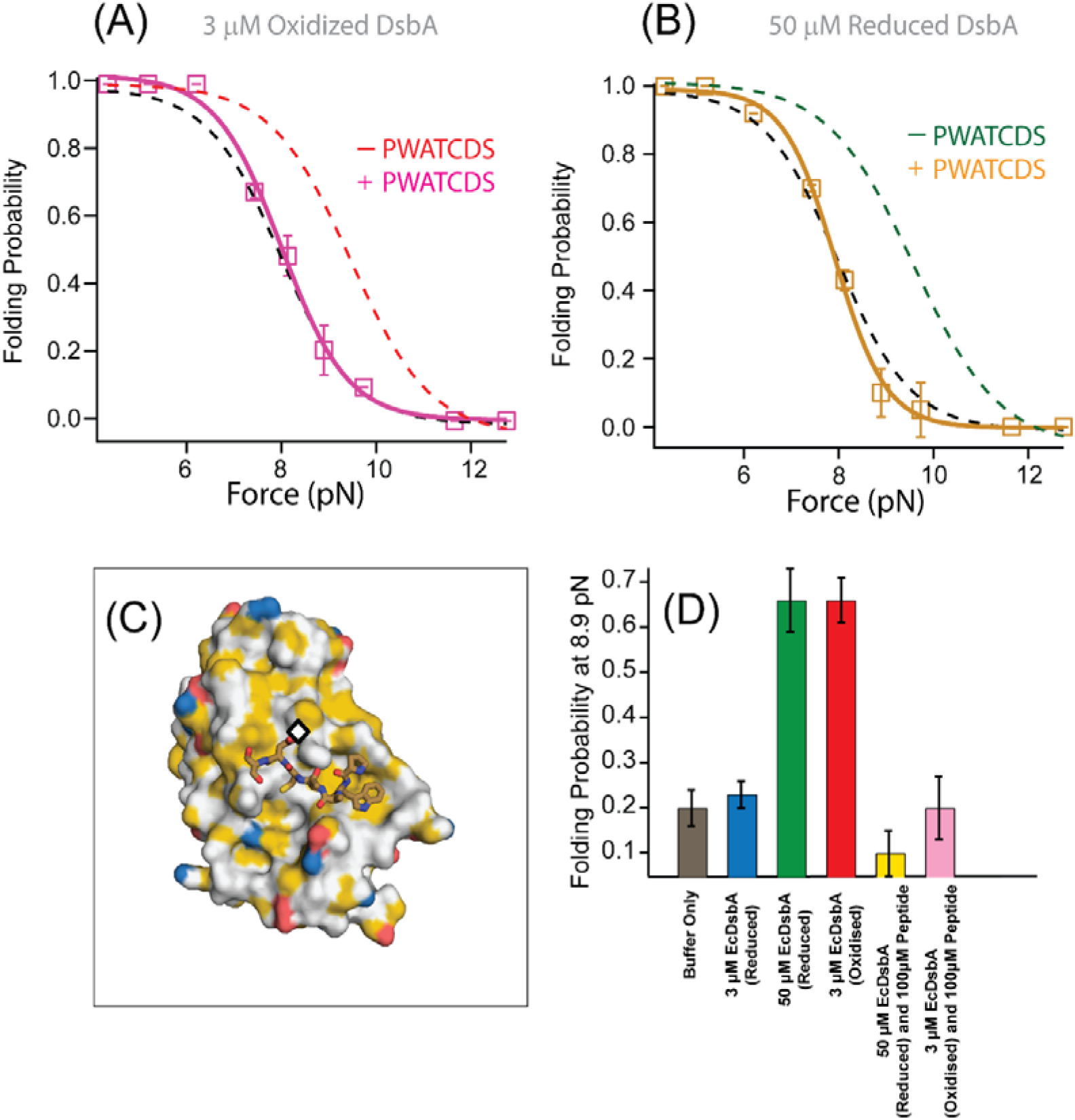
Peptide inhibitor of DsbA maps the chaperone activity to the groove surrounding the catalytic site. **(A)** The folding probability of protein L in the presence of 3 μM oxidized DsbA and 100 μM PWATCDS peptide (pink squares) is shifted back to the folding probability of the control experiment (no DsbA or peptide, black dotted line), indicating inhibition of the mechanical chaperone activity of oxidized DsbA (red dotted line fittings). More than five individual trajectories are measured and averaged for each force. Error bars are standard errors of mean **(B)** The mechanical chaperone activity of 50 μM reduced DsbA is also inhibited in the presence of 100 μM peptide (gold squares), aligning with the folding probability in the absence of DsbA (black dotted line fittings). More than five individual trajectories are measured and averaged for each force. Error bars are standard errors of mean **(C)** Structure of *Proteus mirabilis* DsbA with PWATCDS peptide bound. The active site *CXXC* motif is marked with a black diamond. The surface surrounding the active site is largely hydrophobic, as colored according to the schema of Hagemans *et al.* whereby any carbon not bound to a heteroatom is colored yellow. **(D)** The lower insert shows the folding probability at 8.9 pN in absence (gray) and presence of 3 μM reduced DsbA (blue), 50 μM reduced DsbA (green), 3 μM oxidized DsbA (red), 50 μM reduced DsbA with 100 μM PWATCDS peptide (yellow) and 3 μM oxidized DsbA with 100 μM PWATCDS peptide (pink).

We repeated the mechanical foldase assay with protein L and either reduced or oxidized DsbA using magnetic tweezers, now in the presence of the PWATCDS peptide. With 3 μM oxidized DsbA at 8.9 pN, the folding probability of protein L is found to be 0.66, however, with the addition of the peptide at 100 μM, the folding probability shifts to 0.21, suggesting a loss of the enzyme’s foldase activity (Fig 4A, pink curve and Supp fig 4). Furthermore, the inhibitory effect is evident at all forces tested (4 – 12 pN), as the folding probability of protein L in the presence of oxidized DsbA and 100 μM peptide closely tracks the folding probability of the control experiment. The residual foldase activity with 50 μM reduced DsbA is also lost upon addition of 100 μM PWATCDS peptide (Fig 4B, gold curve and Supp Fig 5). A comparison at the highly sensitive equilibrium force of 8.9 pN demonstrates the large shifts in the folding probability of protein L with or without the catalytic thiols oxidized versus in the presence or absence of the PWATCDS peptide (Fig 4D).

### Equilibrium folding energy calculation of protein L in the presence and absence of DsbA

Equilibrium folding energy of protein L has been determined as a function of force, based on Boltzmann distribution, using the following equation, as described by Chen et al (65).

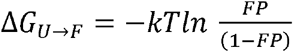

Supplementary Figure 6 shows a comparative data of free energy difference (ΔG) within a span of 6-10 pN force, where both folding-unfolding transitions occur within experimental time-scale. Our result shows, at a given force, the free energy of folding of proteinL, in the presence of oxidized DsbA, is much less than in absence of DsbA (Supplementary Figure 6A). This clearly shows the foldase nature of oxidized DsbA. However, in presence of the reduced DsbA, at 3uM, the free energy of folding is almost same with that of the control. But, in the presence of 50uM reduced DsbA, the free energy of folding decreases, which indicate that the energy landscape shifted towards to folded state (Supplementary Figure 6B). Addition of peptide to the 3μM oxidized or 50μM reduced DsbA decrease the folding as both the cases the free energy of folding is higher in comparison to without peptides (Supplementary Figure 6C and Supplementary Figure 6D).

## Discussion

By applying single-molecule force spectroscopy using a novel magnetic tweezers assay, we have provided an empiric description and measurement of a redox-driven chaperone behavior of *E. coli* DsbA. We use a protein domain that lacks cysteines to show that the chaperone activity of DsbA is separable from its oxidoreductase activity. Based on our observations, we attribute the acceleration of folding to transient noncovalent interactions between DsbA and its substrate that do not involve thiol/disulfide exchange with said substrate. We have further been able to localize this interface to the hydrophobic groove surrounding the *CXXC* catalytic motif by taking advantage of the reactivity of the catalytic thiols, as well as the groove affinity for its DsbB electron shuttling partner. Reducing the catalytic thiols or binding of the PWATCDS peptide both alter the interface between DsbA and the substrate protein L such that chaperone activity is weakened or abrogated Determining if these two modifications of the DsbA enzyme alter electrostatics, hydrophobicity, or local geometry would help to elucidate the mechanism by which DsbA recognizes and folds its protein L substrate (45). Interestingly, conformational changes of DsbA upon oxidation/reduction of its disulfide bond have been noted previously (40, 41), and may contribute to the results seen in this study. Although this is the first report of redox-controlled mechanical foldase activity for DsbA, there are several other instances of chaperones with redox activated switches. Such a mechanism can be rationalized as a protective response to increased oxidative stress, which is known to alter side chain chemistry and cause an accumulation of misfolded proteins. Peroxiredoxins act as oxidoreductases in monomeric form but oligomerize into molecular chaperones under conditions of oxidative stress (46, 47). S-nitrosylation, a by-product of reactive nitrogen stress, at a non-catalytic thiol of human thioredoxin regulates its redox and anti-apoptotic activity (48).

Foldase activity of DsbA could have broader implications for periplasmic protein quality control. DsbA has been shown, for example, to bind to translocating polypeptides in the periplasm, both co-translationally and post-translationally(*1*, *49*). Here we propose an alternative version of chaperone biased polypeptide translocation based on our experimentally observed results that could be important for post-translationally secreted polypeptides (Fig 5). We suggest that DsbA enhanced folding of protein domains on the periplasmic mouth of the Sec pore can generate a pulling force that transfers its strain to the polypeptide in the translocon tunnel, and to any portion still in the cytosol. Chaperone assisted folding on the periplasmic side of the membrane would then in turn decrease the energetic costs of protein translocation.

**Figure 5:**
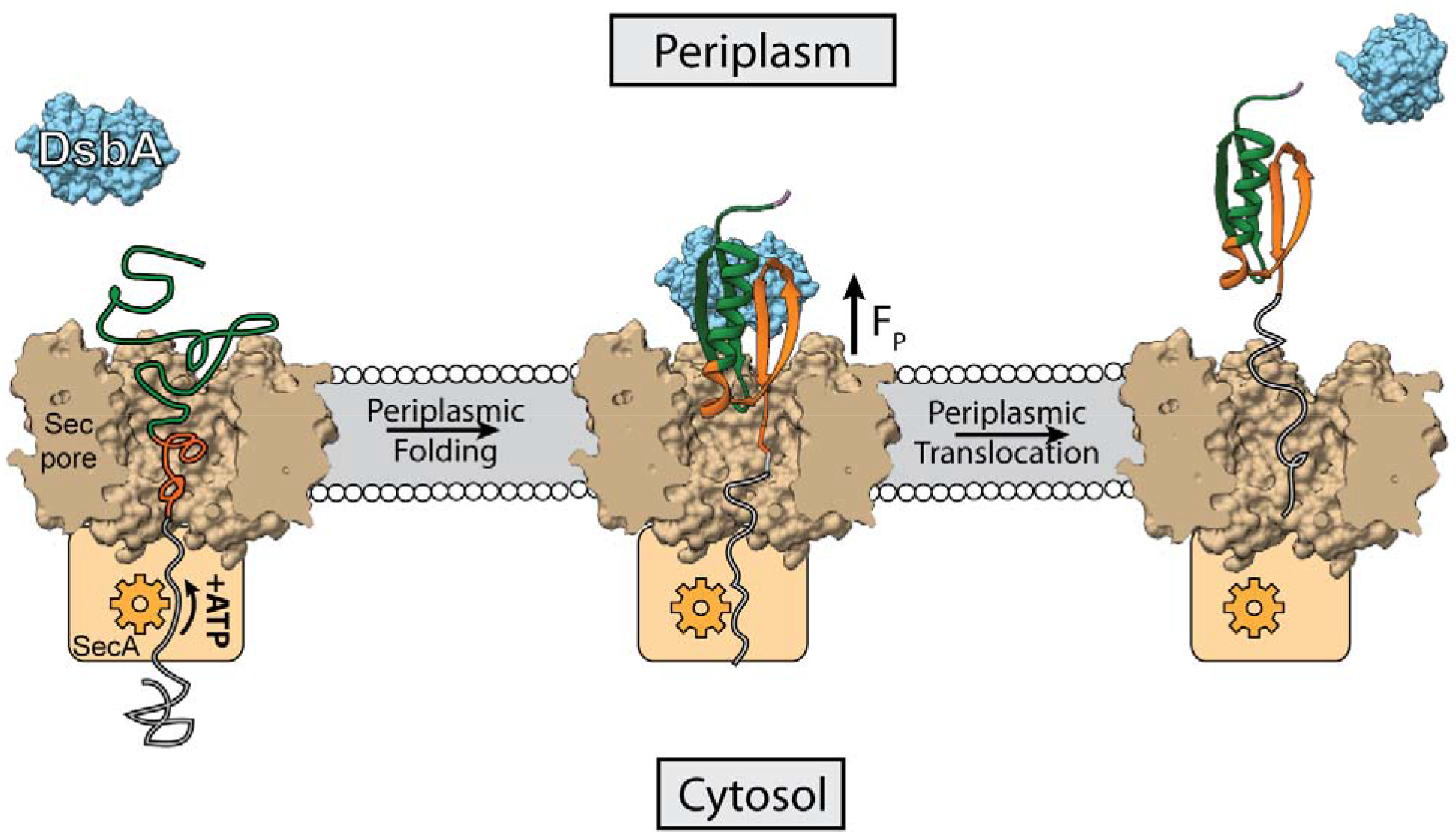
DsbA accelerated transport across the translocon pore. Translocation of peptides into the periplasm take place with the help of Sec machinery, where SecA (orange) translocates the peptide with the help of ATP hydrolysis. Here we show the portion of the translocated polypeptide that is exposed to the periplasm (green polymer) may interact with oxidized DsbA (light blue) and fold on the mouth of the translocon pore, generating a pulling force (F_P_) of several piconewtons. The pulling force strains the remaining polypeptide in the translocon pore (grey polypeptide) and overcoming the friction of the tunnel, pulls it through to the periplasmic side. The work of protein folding therefore reduces the number of rounds of ATP hydrolysis that the SecA motor must undergo to push the polypeptide through the translocon pore.

Prior to translocation, a protein is first maintained in the unfolded state by the secB chaperone which carries the unfolded polypeptide to the SecA motor for transport of the polypeptide through the SecYEG pore using ATP hydrolysis(*50*). It has been shown for the SecA motor that one round of ATP binding and hydrolysis is responsible for the translocation of 20 amino acid residues through the translocon pore(*51*), so a single protein L domain with 60 residues participating in the fold would require 3 ATP molecules for translocation. Assuming a mechanochemical coupling efficiency of ~50% for the SecA motor(*52*, *53*), hydrolysis of 3 ATP molecules would generate 150 zJ of mechanical work (100 zJ per ATP * 50% efficiency * 3 molecules ATP). Although there are no single molecule force spectroscopy measurements of the forces generated by the SecA motor, it is thought to operate optimally over the force range of 5 – 11 pN, stalling at the higher forces, which holds true for most protein translocating motors(*54*). This range coincides with the forces over which DsbA can assist protein folding, reaching a maximal effect at 9 pN (Fig 3B). Refolding of a single protein L domain on the periplasmic side at 9 pN with a step size of 9.5 nm potentially generates 85 zJ (9 pN * 9.5 nm) of work, although the average amount of work performed is determined by the folding probability. This work of protein folding supplies a pulling force from the periplasm that can be harnessed to lower the number of ATP consumed by SecA for translocation. Without DsbA in the periplasm, folding of protein L can only generate 17.1 zJ of energy (9 pN * 9.5 nm *0.2 folding probability - see Figure 3B). However, the presence of DsbA increases the work done by folding to 56 zJ (9 pN * 9.5 nm * 0.66 folding probability). Thus the work of protein folding, assisted by the DsbA chaperone can supply one-third of the energy needed for protein L translocation, lowering the ATP consumption to only 2 molecules per protein L polypeptide translocated. If the SecA motor has even lower efficiency, as suggested by certain studies demonstrating the requirement of 1,000 ATP molecules per proOmpA transported, then the mechanical work done by DsbA is of even greater importance(*54*).

There is emerging evidence to suggest that many chaperones transmit mechanical forces to their substrates during folding. Trigger Factor increases the transmission force of the polypeptide through the ribosomal tunnel by assisting protein folding at higher forces(*26*, *55*). The results from the present study show how the chaperone activity of DsbA could assist the protein to translocate from the cytoplasmic end to the periplasmic end. Besides DsbA, other chaperones like HSP70 and BiP are known to assist the translocation process(*15*, *16*). It has even been demonstrated that high stability titin domains are transported into the lumen of the mitochondria with the help of mtHSP70(*17*). Another common chaperonin, GroEL, mechanically unfolds the misfolded proteins on its apical domain(*56*). Thus, a similar mechanical chaperone activity by all these chaperones indicates a broad mechanical role of chaperones which might be required in many biological process such as rescuing of stalled polypeptide chains in the translation process, accelerating protein folding in elastic tissues, or biasing translocation of polypeptides from one cellular compartment to another.

As a factor involved in the maturation of a diverse set of bacterial exotoxins, adhesins, and secretion machinery, DsbA affords a single attractive target for attenuating a range of virulence phenotypes(*9*, *43*). Deletions of DsbA have been shown to alter the course of infection in animal models of *E. coli*(*11*) and *P. mirabilis* urinary tract infection(*57*), and *F. tularensis* bacteremia(*58*), among others. Moreover, the loss of DsbA activity does not appreciably alter cell viability, lessening the selective drive for antibiotic resistance(*59*). To date, anti-DsbA therapeutic development has largely focused on inhibiting its oxidoreductase activity localized to the catalytic hydrophobic groove(*8*, *9*, *60*). Recent efforts have also explored potential small molecule and peptide binders of the non-catalytic groove on the opposite face, thought to be involved in protein-protein interactions(*10*, *61*). In this context, it is noteworthy that the PWATCDS peptide which binds at the catalytic hydrophobic groove and inhibits the DsbA oxidase also inhibits the mechanical foldase. In a related drug development program, a class of small molecules termed ‘pilicides’ are capable of preventing pilus formation by disrupting chaperone-mediated folding and assembly of the Chaperone-Usher system(*62*–*64*). Along a similar vein, we propose that the redox-controlled DsbA chaperone activities afford a novel avenue for anti-virulence antibiotic development.

## Methods

### Protein expression and purification

DsbA was purified from *E. coli* as described previously(*34*). The DsbA was used fresh after purification with storage at 4C and never after freezing. Oxidized DsbA was generated by incubating the enzyme in 10 mM oxidized glutathione overnight at 4° C. Reduced DsbA was made by incubation with 100 μM TCEP overnight. The GSSG or TCEP was then removed by 3× 1,000 fold dilution into phosphate buffered saline (PBS) and centrifugal concentration. The oxidation state and concentration of the DsbA was determined with a DTNB assay and absorbance at 412 nm and 280 nm respectively using the Ellman’s test protocol described in Goldbio protocol (https://www.goldbio.com/documents/2359/Ellmans+Test+Protocol.pdf). The eight-repeat protein L construct along with AviTag at C-terminus and HaloTag enzyme at N-terminus was expressed and purified as described before(*27*). I27^C32-C75^ was expressed and purified as previously described(*25*, *33*). PWATCDS peptide (99.3%) was purchased from Biomatik. Experiments with the PWATCDS peptide contained 10 uM TCEP to prevent oxidation of the peptide thiols. Paramagnetic Dynabeads (M-270) coated with streptavidin were purchased from Invitrogen. All the experiments were done using PBS as buffer.

### Magnetic tweezers instrumentation and coverslip preparation

Magnetic tweezers experiments were accomplished in our custom-made magnetic tweezers setups, where an inverted microscope (Zeiss Axiovert S100) mounted on a nanofocusing piezo actuator (P-725 PIFOC, Physik Intrumente), a magnet-positioning voice coil (LFA-2010, Equipment Solutions), and a high-speed camera (xiQ CMOS, Ximea GmbH). Fluid chambers are made of two cover slips sandwiched by two strips of parafilm. Both the top and bottom cover slips were washed by sonicating them separately for 30 min in 1% Hellmanex solution, 30 min in acetone, and 30 min in ethanol. The bottom slides are silanized by dipping them in a solution of (3-aminopropyl)-trimethoxysilane (Sigma-Aldrich) 0.1% v/v for 30 minutes in ethanol. After salinization, the bottom glasses are then dried at 100° C for an hour and then sandwiched with washed coverslips using laser-cut strips of parafilm, with a gentle heating at 85° C for 1 minutes. Then the chamber was functionalized with glutaraldehyde, nonmagnetic polystyrene beads (3 μm) and HaloTag amine ligand respectively as described previously(*27*). Recombinant octamer protein was tethered using HaloTag chemistry (N-terminal) and the other end of the protein was decorated with a single paramagnetic bead through biotin-streptavidin chemistry (AviTag, Avidity Technology). Position of the permanent magnets, was controlled with a linear voice-coil with a speed of ~0.7 m/s speed and 150 nm position resolution. The force on the protein is calibrated by the procedure mentioned by Popa et al (27). Experiments with DsbA are performed after >15 minute equilibration with Protein L in the fluid chamber. Similarly, PWATCDS peptide are incubated for > 30 minutes with DsbA before adding to the fluid chamber.

### Data analysis

Only proteins showing eight unfolding events in the high force (45 pN) fingerprint pulse are considered for analysis. After unfolding the protein, the force is relaxed during the “quench” pulse to achieve an equilibrium between the folding and unfolding states to calculate the folding probability. We measure this by calculating the probability of finding the system in a particular state, considering that *t_i_* is the total time the system spends in state *i* (where *i* equals the number of folded domains, *i* = 0, 1, …, *N*) and *t_t_* the total time of experiment, *t_t_* = *Σ_i_ t_i_*. Then, the folding probability is *P_f_* = *Σ_i_ iπ_i_*/*N*, where *π_i_* = *t_i_/t_t_*. This was the same metric used in previous magnetic tweezers based chaperone assays(*26*).

## Acknowledgement

We sincerely thank Prof Julio Fernandez for his kind help in this project. SH thank Ashoka University for support and funding. SH thanks DST core research grant (SERB) and Ramalingaswami Re-entry Fellowship, DBT for funding. We would like to acknowledge Thomas Kahn, Andres Rivas-Pardo and Guillermo Alvarez de Toledo for the preliminary studies on DsbA with protein L.

## Author Contributions

S.H., E.C.E., and D.J.E. designed the project, S.H., E.C.E., D.C., and S.C. performed the experiments and analyzed the data, S.H., D.J.E., and E.C.E. wrote the paper.

## Supplemental Information

**Supplementary Figure 1.**
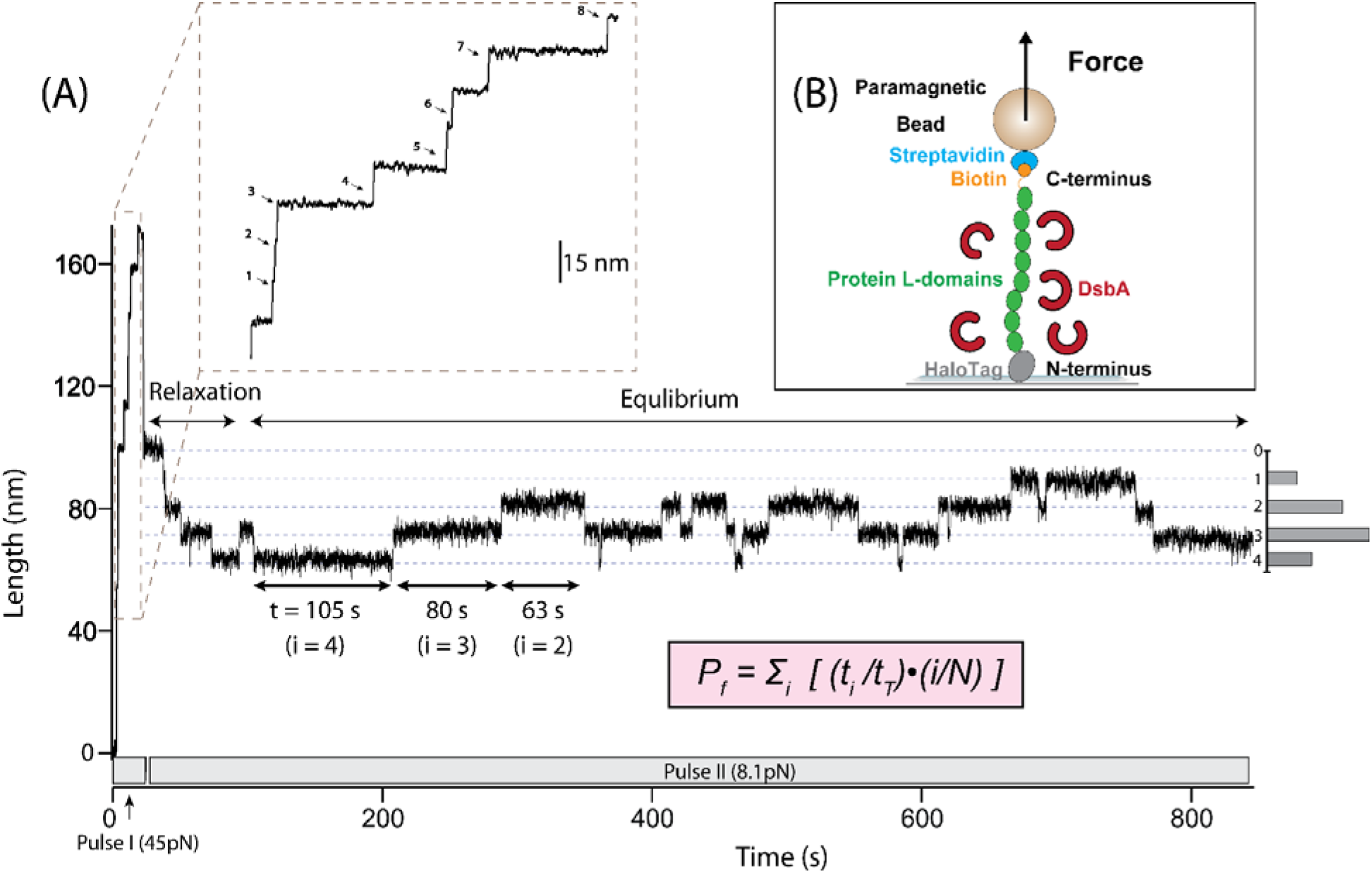
Folding probability under force measured by magnetic tweezers (A) Representative magnetic tweezers demonstrating unfolding and refolding transitions of an eight-repeat (*N*=8) construct of the protein L domain: The protein is first unfolded at a constant force of 45 pN (Pulse I) resulting in eight consecutive unfolding (upwards-step) transitions of 15 nm each (see inset for magnified). The force is then quenched to 8.1 pN (Pulse II) resulting in entropic recoil of the protein followed by relaxation to an equilibrium between folding (downwards-step) and unfolding transitions. The state *i* of the protein at any point during the recording is equal to the number of folded domains. The total residence time (*t_i_*) of each folded state during the equilibrium phase is shown by the gray histogram on the right and the sum of all residence times (*t_T_*) is equal to the duration of the equilibrium phase. After calculating the states *i* and the residence times *t_i_* the folding probability at a force of 8.1 pN is calculated according the presented equation (pink inset). **(B) Schematic of the magnetic tweezers experiment:** One end of octamer of protein L is attached to the glass coverslip via HaloTag covalent chemistry and the other end is tethered to a paramagnetic bead via biotin-streptavidin binding. A precise pulling force is applied by positioning a pair of permanent magnets above the tethered paramagnetic bead with sub-micron resolution. DsbA (red curls) can be washed into or out of the flow cell during the course of an experiment.

**Supplementary Figure 2:**
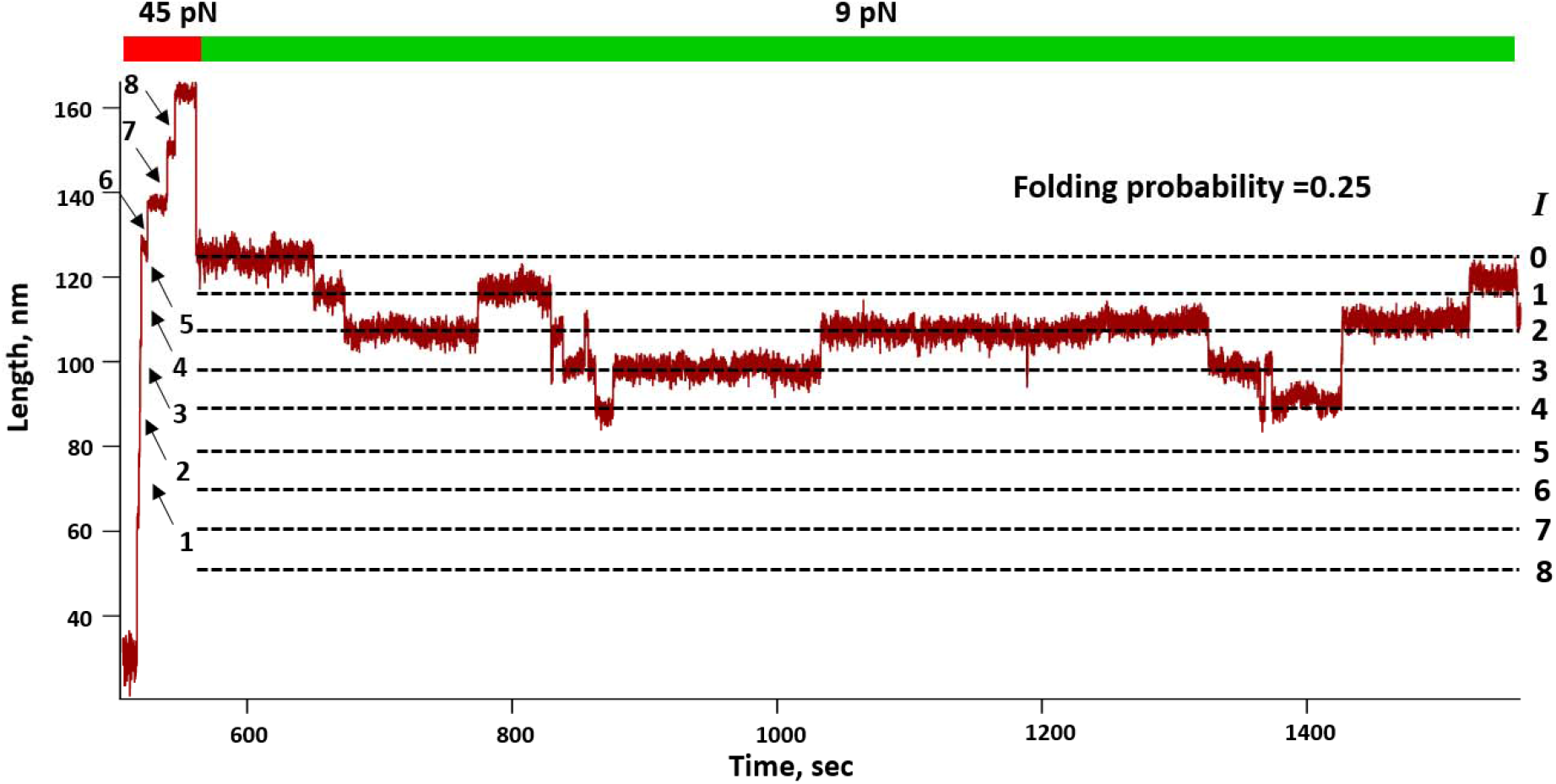
Representative trace of protein L in the presence of 3μM reduced DsbA: In the first pulse, protein L is completely unfolded at 45 pN, where it shows eight unfolding (finger print) steps. Then in the next pulse, it is quenched to 9 pN, where proteins stays in an equilibrium state and showing folding-unfolding dynamics. The calculated folding probability at 9 pN is 0.25.

**Supplementary Figure 3:**
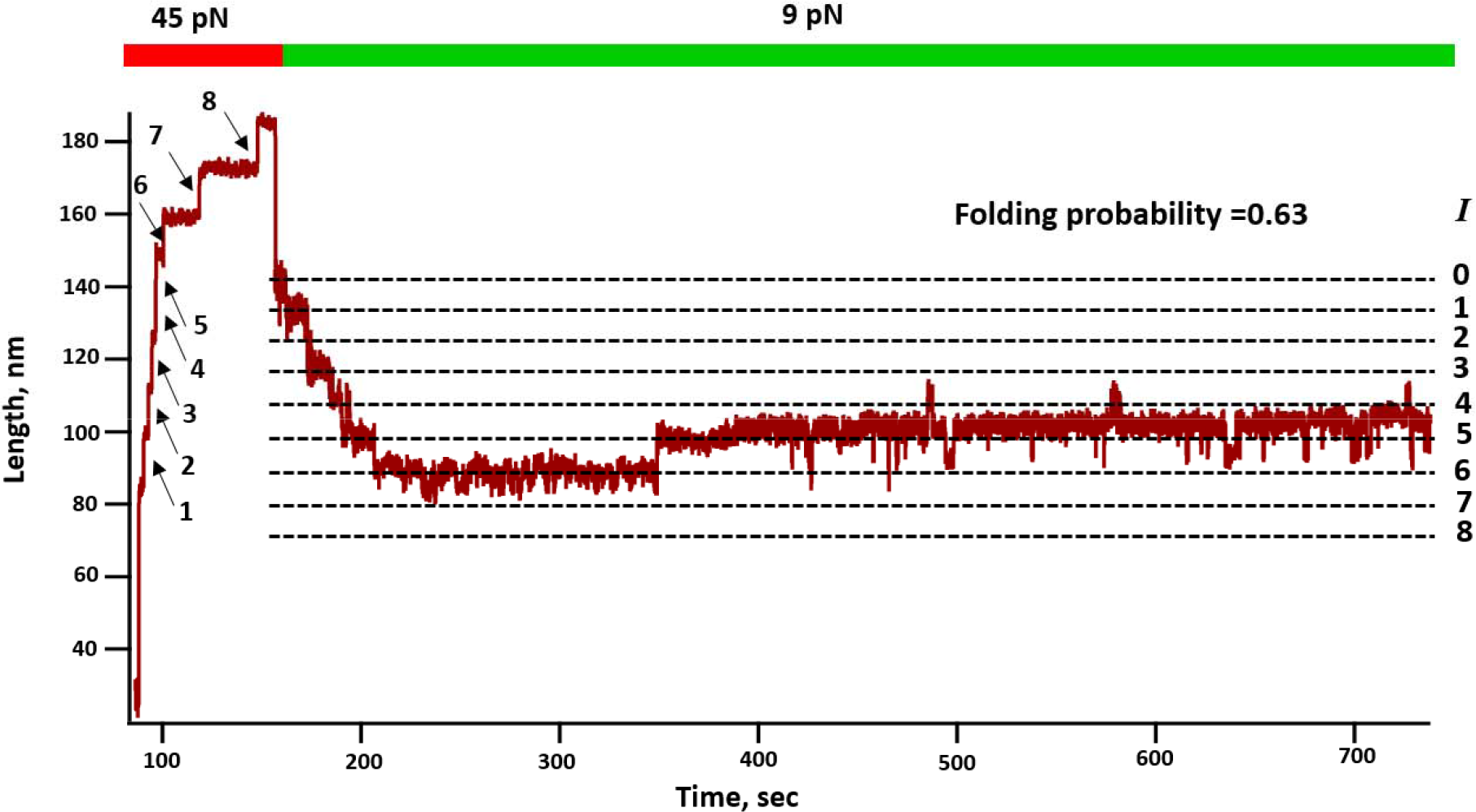
Representative trace of protein L in the presence of 50 μM reduced DsbA: Similar to supplementary Figure 1, in the first pulse protein L is unfolded at 45pN and then quenched at pN. In presence of 50 μM reduced DsbA, folding probability protein L increased to 0.63 at 9 pN.

**Supplementary Figure 4:**
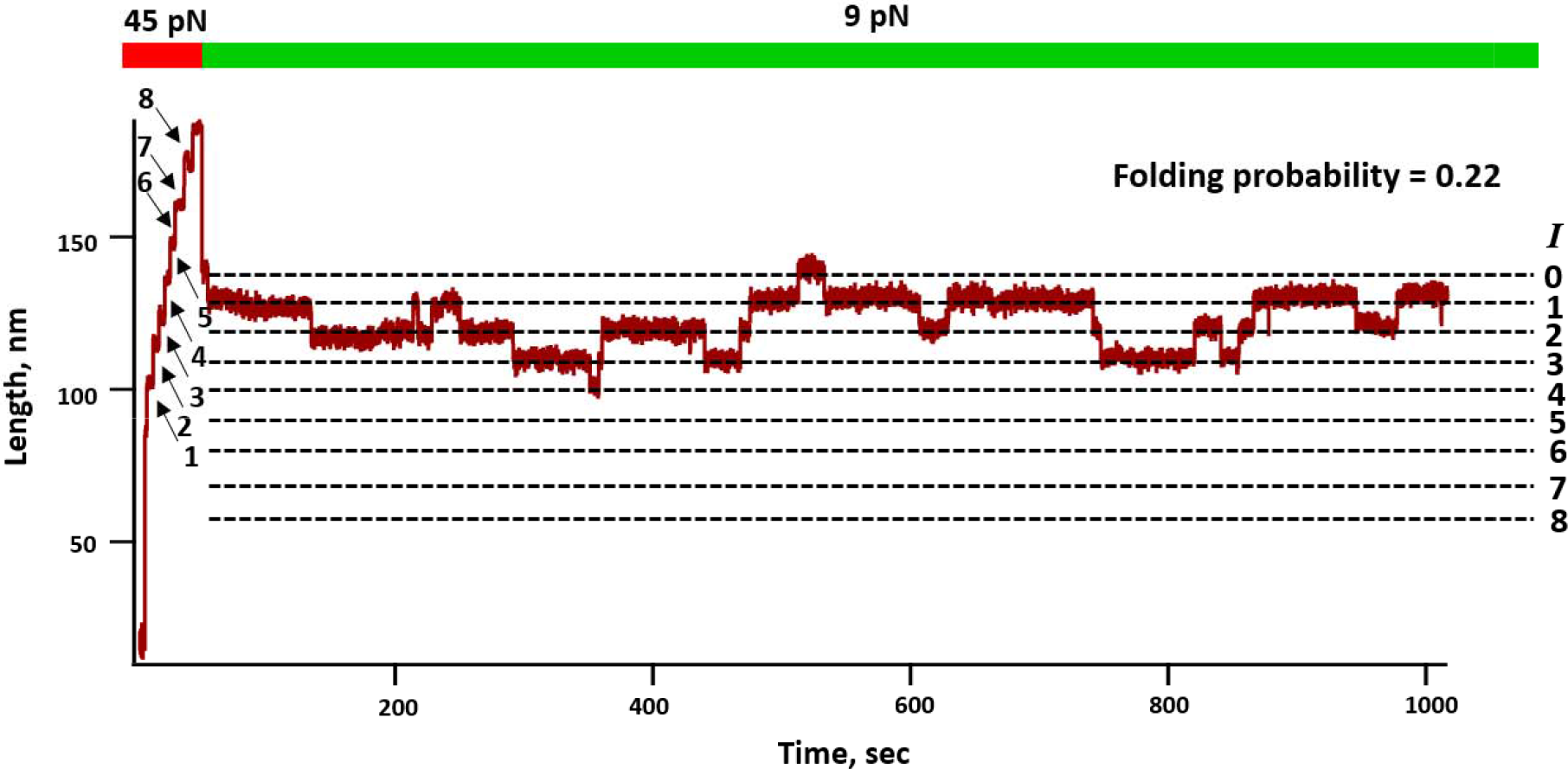
Representative trace of protein L in presence of 3 μM DsbA and 100 μM PWATCDS peptide: Similar to first two figures, protein L was unfolded and folded in the presence of 3 μM oxidised DsbA and 100 μM PWATCDS peptide. In presence of the peptide, the folding probability of Protein L decreases to 0.22 at 9 pN.

**Supplementary Figure 5:**
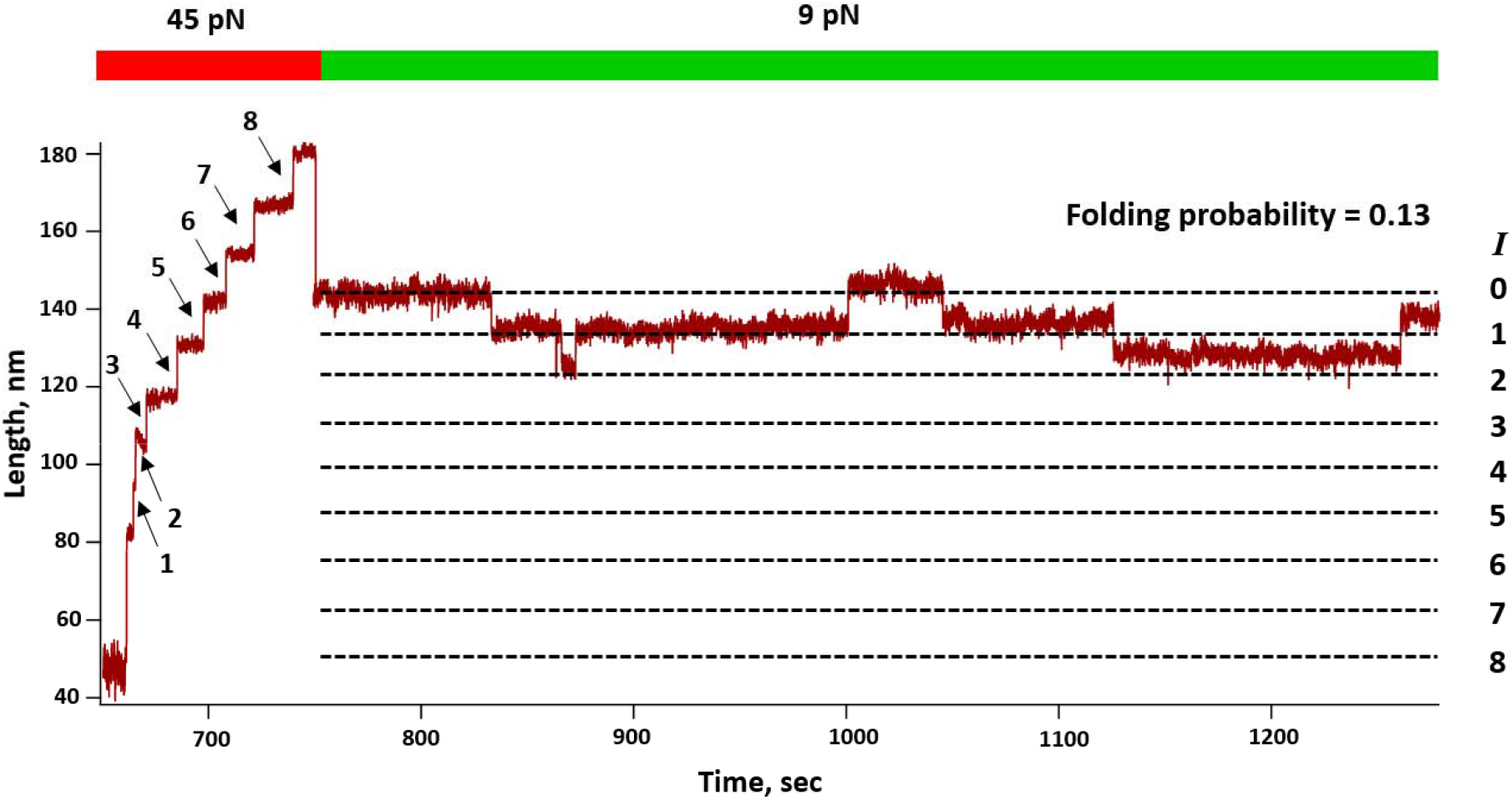
Representative trace of protein L in the presence of 50 μM reduced DsbA and 100 μM peptide: Similar to previous 3 figures, in the first pulse the protein unfolds fully at 45 pN and then in the second pulse, it refolds to 9 pN force. In presence of peptide protein L shows a folding probability of 0.13.

**Supplementary Figure 6:**
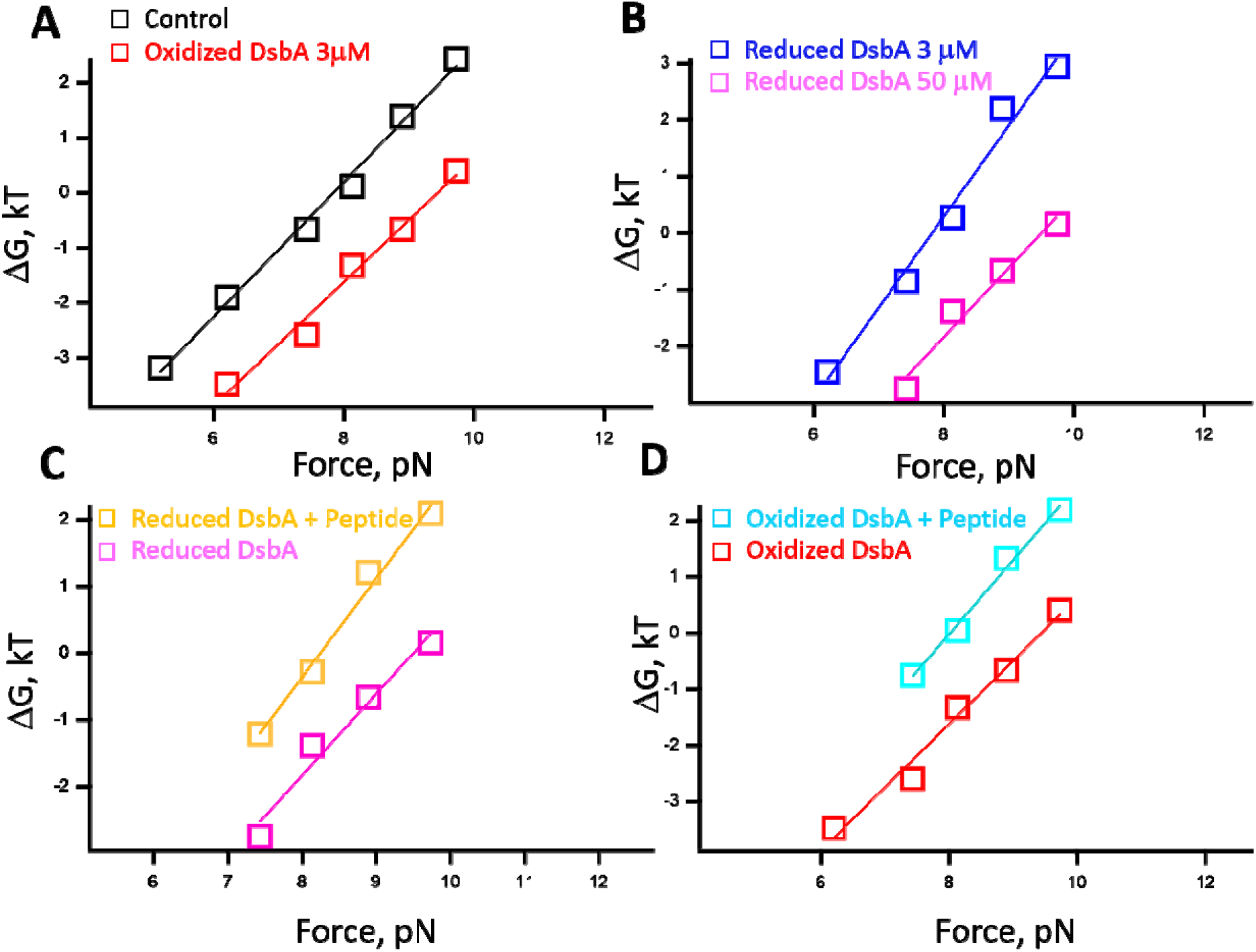
ΔG folding is calculated from the folding probability values using the protocol described in Chen et al., (JACS, 2015, 137 (10), 3540-3546). The ΔG folding is measure for protein L in the presence of 3 μM oxidized DsbA (red), 3 μM reduced DsbA (blue), 50 μM reduced DsbA (pink), 3 μM oxidized DsbA and peptide (cyan), 50 μM reduced DsbA and peptide (yellow) and control (black).

## MFPT Calculations

Mean first passage times (MFPT) were calculated by averaging the trajectory durations for n > 5 molecules. To calculate the folding MFPT, the protein construct was first fully unfolded, the force was relaxed and the time required to achieve folding of all eight domains simultaneously was recorded. This was performed for several different folding forces in the range of 4-8 pN and was then fit with a single exponential equation of the form, and the fit parameters are recorded below:

**Supplementary Table 1:** MFPT refolding fit parameters for DsbA and control conditions with protein L. Likewise for MFPT of unfolding, the protein construct is first allowed to fold completely at a low force before it is probed at a high force of 30-50 pN. The time required to pass to the state with all eight domains unfolded simultaneously is recorded and averaged for several molecules. The same equation is used to fit the MFPT and the parameters are recorded below:

| | $A$ (ms) | $\phi$ (pN) |
| --- | --- | --- |
| DsbA | 0.784 | -0.632 |
| Control | 0.269 | -0.505 |

**Supplementary Table 2:**
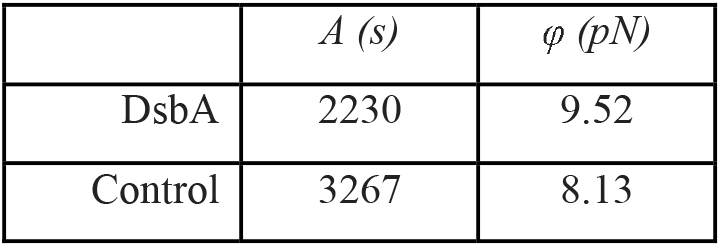
MFPT unfolding fit parameters for DsbA and control conditions with protein L.

